# 3D imaging reveals apical stem cell responses to ambient temperature

**DOI:** 10.1101/2023.05.04.539361

**Authors:** Christian Wenzl, Jan U. Lohmann

## Abstract

Plant growth is driven by apical meristems at the shoot and root growth points, which comprise continuously active stem cell populations. While many of the key factors involved in homeostasis of the shoot apical meristem (SAM) have been extensively studied under artificial constant growth conditions, only little is known how variations in the environment affect the underlying regulatory network. To shed light on the responses of the SAM to ambient temperature, we combined 3D live imaging of fluorescent reporter lines that allowed us to monitor the activity of two key regulators of stem cell homeostasis in the SAM namely *CLAVATA3* (*CLV3)* and *WUSCHEL (WUS),* with computational image analysis to derive morphological and cellular parameters of the SAM. Whereas *CLV3* expression marks the stem cell population, *WUS* promoter activity is confined to the organizing center (OC), the niche cells adjacent to the stem cells, hence allowing us to record on the two central cell populations of the SAM. Applying an integrated computational analysis of our data we found that variations in ambient temperature not only led to specific changes in spatial expression patterns of key regulators of SAM homeostasis, but also correlated with modifications in overall cellular organization and shoot meristem morphology.

## Introduction

Plants continuously form organs during post-embryonic development while at the same time they permanently adapt to changing environmental conditions. The origin of cells forming most of the aerial organs of a plant is the shoot apical meristem (SAM). Here the coordinated interplay of phytohormone activity, mechanical and biophysical constraints as well as transcriptional regulation and cell to cell communication leads to the formation of new organs with remarkable plasticity. The constant supply of new cells that fuels this lifelong formation of new organs is secured by a population of pluripotent stem cells that resides at the very tip of the SAM in the central zone (CZ). Upon cell division stem cell progenitors are displaced towards the peripheral zone (PZ). Eventually, they will be positioned outside the stem cell domain where they start to differentiate and become incorporated into newly formed organs like leaves or flowers. Stem cell fate is triggered by the activity of a distinct population of cells in the so-called organizing center (OC). The cells of the OC are located right beneath the stem cells of the CZ. They express the mobile homeodomain transcription factor WUS which acts in a non-cell autonomous fashion to induce stem cell fate in the apically located stem cell population (Daum et al., 2014; Mayer et al., 1998; Yadav et al., 2011). In response, stem cells express *CLV3,* which codes for a small secreted peptide that can repress the expression of *WUS* in the cells of the OC via a complex signaling pathway (Brand et al., 2000; Fletcher et al., 1999; Jeong et al., 1999; Müller et al., 2008; Schoof et al., 2000).

While this central regulatory unit has been extensively studied under standard growth conditions it is largely unknown if and how the underlying regulatory network is modulated by environmental cues. Plants are continuously exposed to large variations in their local environment with the availability of water, the intensity and spectral composition of light, as well as ambient temperature being the most important abiotic factors. It has recently been reported that ambient temperature has an impact on shoot regeneration (Lambolez et al., 2022), a process that closely resembles SAM function. The overall response to temperature is called thermomorphogenesis and is characterized by increased hypocotyl growth, elongated petioles, hyponastic leaf growth, reduced number of stomata and early flowering (reviewed in Casal and Balasubramanian, 2019; Quint et al., 2016). These temperature induced phenotypes exhibit similarities to shade avoidance responses and not surprisingly it has been shown that both processes are partially regulated by the same genes. The photoreceptor PhyB has been reported to contribute to temperature sensing via a thermal reversion of its active state Pfr to its inactive form Pr (Jung et al., 2016; Legris et al., 2016). The transcriptional response to changes in ambient temperature is largely mediated by transcription factors PIF4 and PIF7 (Fiorucci et al., 2020; Koini et al., 2009). Active phyB inhibits PIF4 activity and prevents PIF4 from binding to its target promoters. At high temperatures increased thermal reversion will lead to increased amounts of the inactive form Pr, resulting in higher *PIF4* mRNA expression which ultimately leads to the expression of auxin biosynthesis genes, growth-promoting factors and genes involved in brassinosteroid biosynthesis. In addition, temperature sensing also depends on *ARP6* a subunit of the conserved *SWR1* complex that is involved in positioning the Histone variant *H2A.Z* into nucleosomes that can modulate transcription in a temperature-dependent manner (Kumar and Wigge, 2010).

Here we developed a specific image analysis workflow to investigate how the stem cell system as well as the entire SAM responds to different temperature conditions. We find that elevated ambient temperature led to an apical shift in *CLV3* as well as *WUS* reporter activity. At the same time, we could observe changes in cell numbers that affected predominantly the L2 layer indicating a layer specific adaptation of cell division rates. By performing temperature shift experiments we describe the temporal dynamics of this response. We also show that the spatial distribution of a WUS-GFP fusion protein is only slightly affected by temperature changes and it remains unclear if this could be a cause for the observed changes in *CLV3* reporter activity.

## Results

### Considerations for reporter design

Using fluorescent reporter lines permits to monitor dynamic changes in gene activity in live tissues. Since the shoot meristem is a complex three-dimensional tissue we wanted to develop tools that allow monitoring of a number of morphological SAM parameters simultaneously. Especially for studying adaptation processes it would be ideal to follow specific cellular processes in the same meristem over an extended period of time during the adaptation period. However, this still presents an enormous challenge in plants due to the relatively slow mode of development and the invasiveness of repeated imaging of the enclosed and protected shoot meristem. It is possible to image SAMs over a course of several days but it is very challenging with low survival rates and small sample numbers. Therefore, we decided to use cohort analysis by doing sacrificial live imaging of meristems from groups of plants that were cultivated under defined growth conditions. While this approach did not allow us to follow the behavior of individual cells, it gave us access to snapshots of meristem development with maximum reproducibility.

We decided to combine up to three transcriptional reporters using different fluorophores fused to a C-terminal nuclear location signal, since this resulted in robust and strong signals and would also facilitate 3D nuclear segmentation. First, we used a ubiquitously expressed reporter *pUBQ10:3xmCherry-NLS:tUBQ10* to be able to analyze a wide range of morphological and cellular parameters of the SAM (Figure1 A,D,E). To gain insight into specific stem cell and niche cell responses we combined this reporter with two cell type specific reporters, namely *pCLV3:mTagBFP2-NLS:tCLV3* and *pCLV3:GFP-NLS:tCLV3, as* well as *pWUS:2xVENUS-NLS:tWUS* (Figure1, B,C,E). Whereas *CLV3* expression marks the stem cell population, *WUS* promoter activity is confined to the organizing center (OC), the niche cells adjacent to the stem cells, hence allowing us to record on the two central cell populations of the SAM. All reporters were constructed using GreenGate cloning (Lampropoulos et al., 2013) and sequentially transformed into *Arabidopsis thaliana Col-0* ecotype Figure 1. The activity of the reporters as analyzed by confocal imaging was in line with previously published expression patterns (Brand et al., 2002; Galvan-Ampudia et al., 2020; Ma et al., 2019). We also created reporter constructs that combined the *CLV3* and *UBQ10* reporter in one T-DNA since it was faster to establish respective lines especially if the transgene was supposed to be introduced in specific mutant backgrounds. The same approach was tested for combinations of the *WUS* and the *CLV3* reporter constructs but we observed a mutual interference on reporter expression, most likely due to the close proximity of regulatory sequences (data not shown).

**Figure 1:**
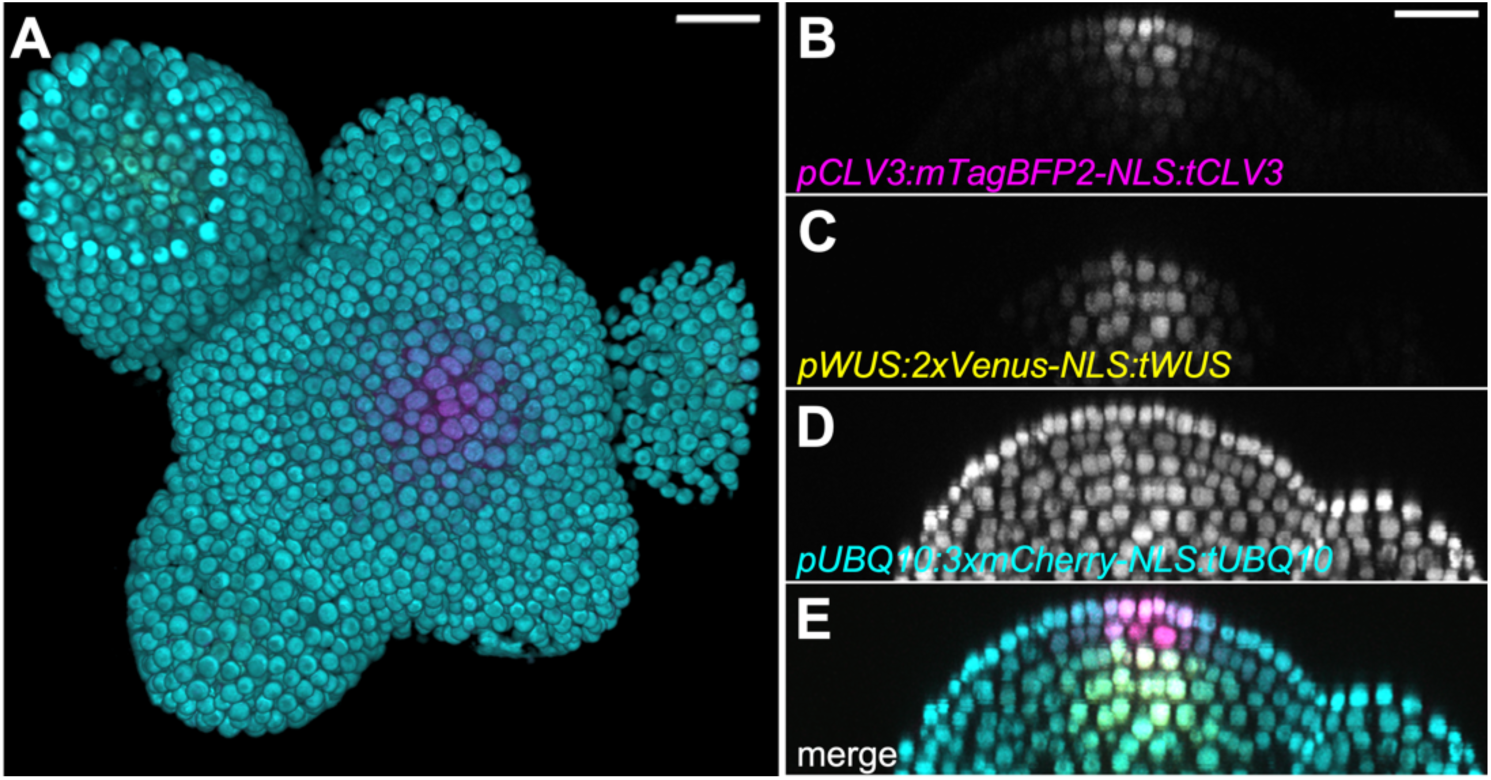
Triple reporter line reveals cellular design of SAM and apical stem cell system. Inflorescence meristem of a plant harboring three different independent reporter constructs was imaged using confocal laser scanning microscopy. A: Top view of a 3D reconstruction of all three reporters, images of all three single channels are shown as yz cross sections. B: *pCLV3:mTagBFP2-NLS:tCLV3*, C: *pWUS:2xVenus-NLS:tWUS*, D: *pUBQ10:3xmCherry-NLS:tUBQ10*. E: merge. Scalebar corresponds to 20 µm.

### Image analysis workflow

The overall workflow of our image analysis pipeline is depicted in Figure 2. Since we wanted to integrate a number of already existing analysis tools to solve different processing tasks we decided to implement our pipeline as a cross platform graphical workflow in KNIME (Berthold et al., 2009). This approach offered several advantages. KNIME is an intuitive and easy to use open-source software platform that allows to integrate a number of commonly used image analysis tools (e.g. Matlab, Fiji, ilastik) into a single workflow thereby assuring seamless data exchange during different processing steps. In addition, it can run analysis workflows fully automatically or with different degrees of user interaction.

**Figure 2:**
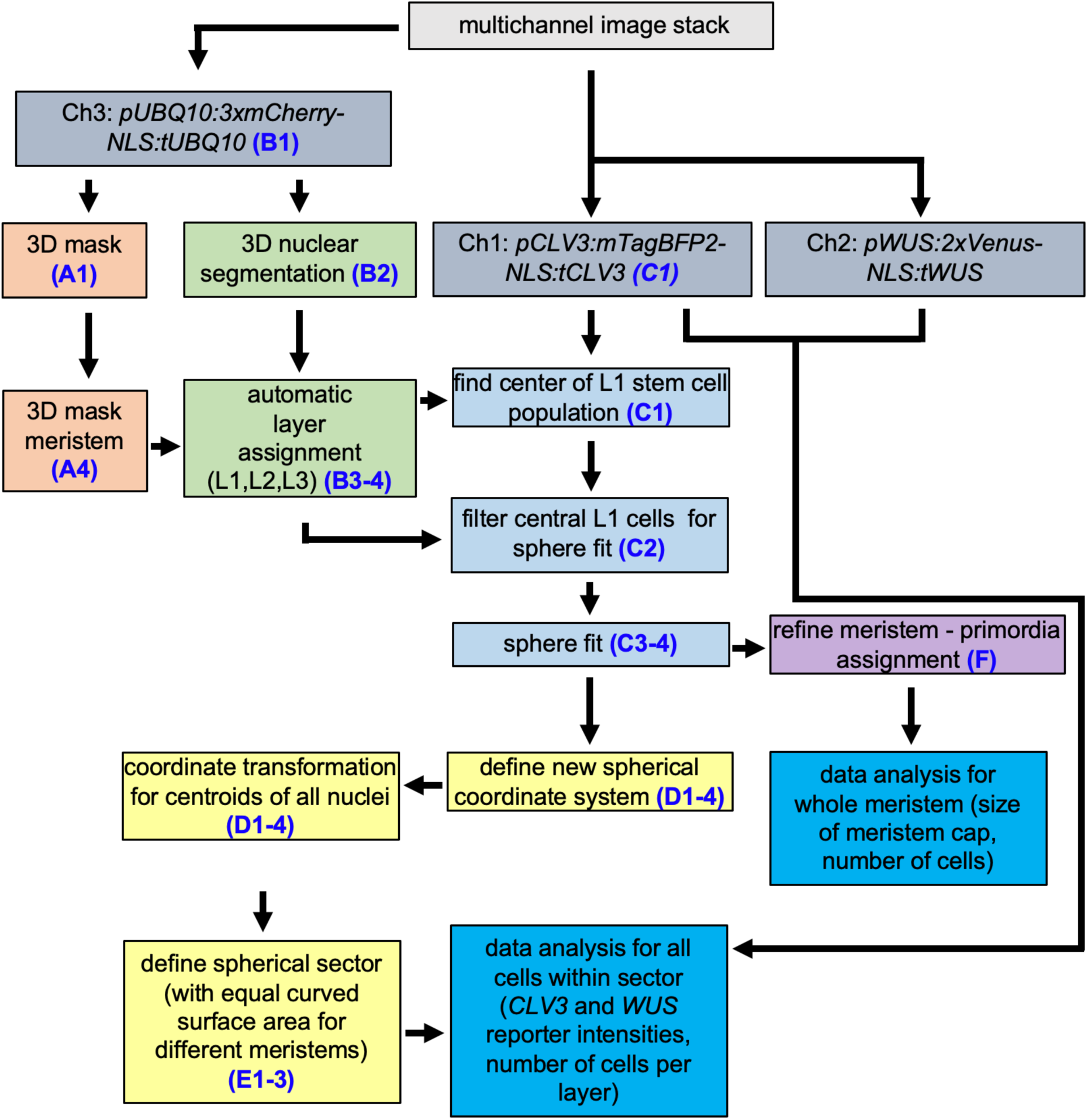
Schematic overview of the main image analysis workflow. Isotropic 3-channel image volumes were the starting point of the main analysis pipeline. The 3D nuclear segmentation step (B2) was run separately on Google Colabs. The rest of the analysis is carried out in KNIME.

The *pUBQ10:3xmCherry-NLS:tUBQ10* channel (Figure3: B1) served as the main backbone to create a 3D binary mask representing the overall shape of the SAM specimen as well as the 3D nuclear segmentation of the entire image volume (Figure3: A1, B2). To obtain a precise nuclear 3D segmentation we used StarDist (Schmidt et al., 2018). Initial training data were obtained by classical 3D seeded watershed segmentation with manual correction of seeding. To our surprise and as outlined in the material and methods section, even 8 pairs of small 3D image-segmentation substacks were sufficient to obtain a trained model that provides robust nuclear segmentations for our imaging conditions. The 3D binary masks were generated after training a pixel classification model in ilastik (Berg et al., 2019). Since expression of the *CLV3* as well as *WUS* reporters was not restricted to the inflorescence meristem but could also be observed in emerging flower primordia, it was necessary to exclude these regions from our analysis. Therefore, the binary masks were spatially restricted by manually drawing separation lines between shoot meristem and primordia (Figure 3: A2). After finding respective connected regions, a binary mask for the analysis of the central part of the inflorescence meristem could be obtained (Figure3: A3,4).

**Figure 3:**
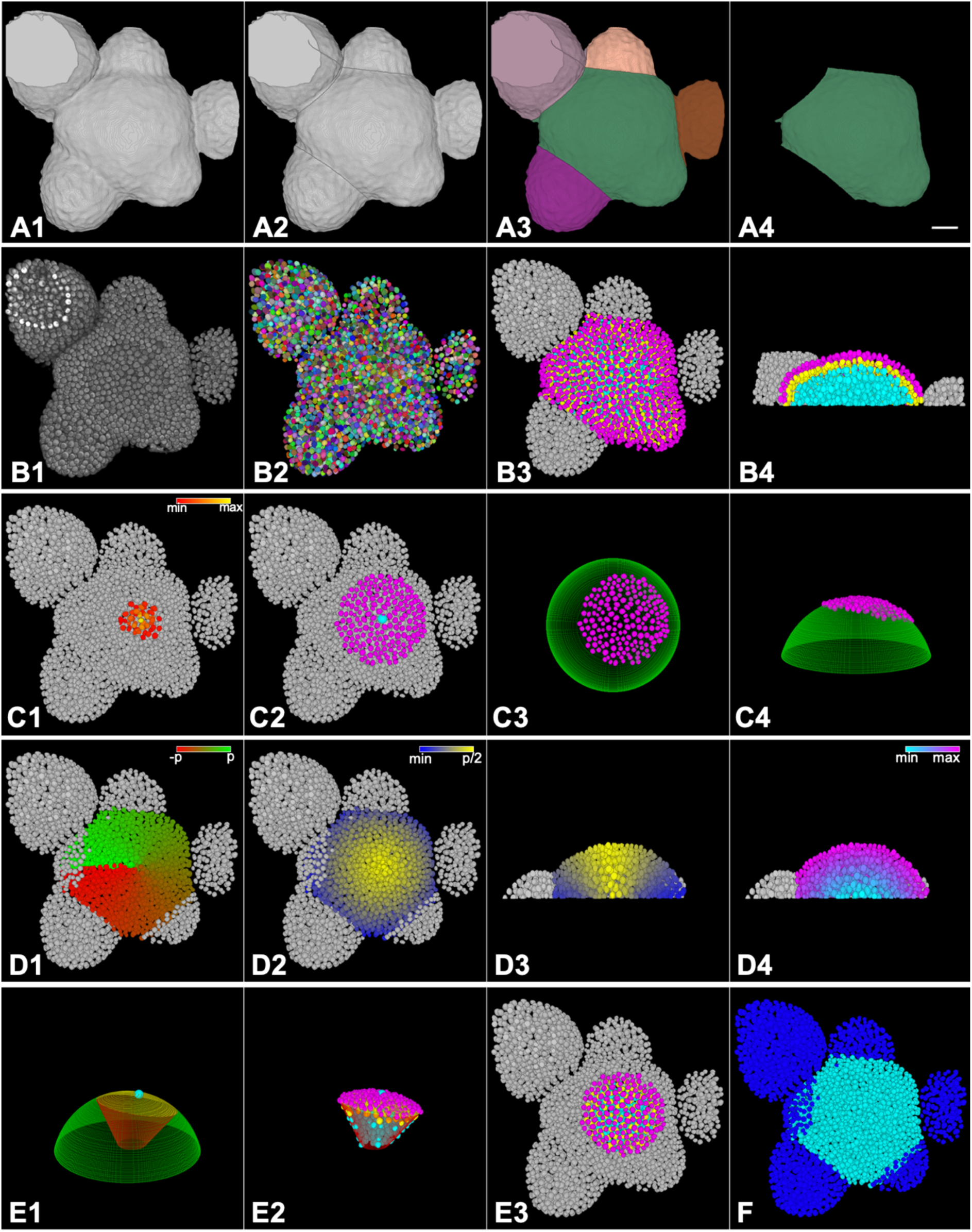
Processing steps of the analysis workflow. Results of key processing steps in our data analysis workflow are visualized for an example stack. Input was a 4D image volume obtained by CLSM of a triple reporter meristem sample. After channel splitting a 3D binary mask (A1-A4) is generated from the *UBQ10* reporter. The raw mask (A1) is further processed so that older primordia are separated from the main meristem by manually drawn lines (A2) followed by a connected component analysis (A3) resulting in a mask that covers only the central meristem and very young primordia (A4). The ubq10 channel (B1) is also used for obtaining a 3D nuclear segmentation (B2) which is combined with the mask generated in (A4) to automatically assign layer identity to all nuclei of the central meristem (B3, B4). B4 shows a y-z view of the meristem in B3. Panel C1-C4 illustrates the process of how epidermal cells within a given distance (can be selected by the user) from the meristem summit point (marked by the center of the *CLV3* expressing stem cells in L1 (C1)) are selected to fit a sphere that can be used to describe the size of the meristem (C2-4). The sphere fit is also the prerequisite for a coordinate transformation from euclidian to spherical. The origin of the new coordinate system is the center point of the sphere with the new z-axis pointing towards the meristem summit point. Corresponding visualizations can be seen for azimuth (D1), elevation (D2 and cross section in D3) and radius (D4). The new coordinate system was used to define a spherical section with a fixed size of the spherically curved surface (yellow region in E1, light blue spot marks the summit point). The quantification of *WUS* and *CLV3* reporter signals was restricted to all nuclei which had their centroids inside the spherical section (E2). The nuclei within the section are visualized as top view in E3. Meristem-Primordia assignment can also be improved by assigning all nuclei inside the sphere to the meristem and all nuclei outside to primordia or other tissue (F). Scalebar corresponds to 20 µm.

One important processing task of our workflow was the automated assignment of segmented nuclei to the clonal cell layers of the SAM. Nuclei were assigned to the outer epidermal cells (L1), the subepidermal cells (L2) or the inner cells of the rib meristem (L3). To achieve this, we calculated the shortest distance of the centroid of each individual nucleus to the surface of the meristem. To this end the 3D binary mask was meshed and the mesh points were used to calculate the distance of the centroid of each nucleus to the closest mesh point on the upper surface of the binary mask of the meristem. For this task we used the Matlab iso2mesh package (Tran et al., 2020) and KNIMEs Matlab integration. Layer assignment can be achieved interactively by user supplied distance thresholds for layers L1 and L2 (Figure3: B3,4).

The size of the shoot apical meristem is often correlated with cytokinin controlled cell proliferation. Cytokinin levels in the SAM are partially regulated by degrading CYTOKININ OXIDASES. For example, SAMs of *ckx3 ckx5* double mutants have significantly enlarged SAMs whereas ectopic expression of *CKX1* causes a reduction in meristem size (Bartrina et al., 2011; Werner et al., 2008). Thus, quantifying meristem size is an important step when comparing SAM activity. The wildtype shoot apical meristem is a dome shaped structure that can be well approximated by a sphere (Ma et al., 2019), which in turn lends itself well for parameter deduction. Hence, in a first step we determined the summit point of the dome shaped SAM by using the signal intensities of the *CLV3* reporter in the L1 stem cells (Figure3: C1 and C2, see materials and methods for details). This was necessary to accommodate for potential tilting of the central meristem axis during mounting of the specimen for imaging. The coordinates of the centroids of L1 cell nuclei located within a user defined distance D_s_ of this summit point were then used to fit a sphere that closely approximates the shape of the SAM (Figure 3: C2-C4). We kept the value for D_s_ constant for most of the analyzed meristems (32,1µm = 100 px). This is possible because the coordinates for the center point of the fitted sphere as well as its radius are stable with just minor variations for D_s_ values higher then 25µm (see Figure S1). However, D_s_ was adapted to accommodate for very small meristem sizes. The radius of the sphere provided a direct quantitative readout for meristem size. However, this method had obvious limitations: If the overall meristem morphology is distorted, such as in *clv3* mutants, a sphere fit may not accurately reflect SAM size. Conversely, sphere fitting also had a number of advantages for downstream processing steps. Firstly, it provided a more unbiased way of meristem size assessment compared to manual measurements. In addition, it allowed the introduction of a coordinate system with the center point of the sphere as origin and the z-axis pointing in the direction of the summit point determined by the L1 *CLV3* reporter maximum. After respective coordinate transformation the position of each nucleus can be easily described using spherical coordinates (Figure3: D1-D4), which in turn allowed us to quantitatively and reproducibly describe morphological variation. For analyzing meristems of plants grown under different conditions we wanted to focus on the most central part of each SAM including the stem cells, the organizing center and the inner cells of the peripheral zone. Introducing a specific SAM coordinate system allowed us to easily define and analyze spherical sections with the stem cell domain at its center (Figure 3 E1-E3). We empirically chose the parameters of these sections based on two criteria: firstly, to ensure comparability the outer surface of the sections had to be of the same size and secondly, we selected only cells that were not incorporated yet in emerging organ primordia and were located within the central dome shaped part of the SAM (see Figure S1). It could be argued that this could lead to quite different ratios of stem cells and transient amplifying cells if very big and very small meristems are to be compared but the size of the stem cell domain is known to scale according to the overall size of the SAM (Gruel et al., 2016). Lastly, the sphere could also be used to further refine the distinction between meristem cells and primordial cells (Figure3: F).

### Ambient temperature influences spatial distribution of stem cells

Since we aimed to challenge the SAM by variations in ambient temperature within the physiological range, we chose to grow *Arabidopsis* plants carrying two reporter genes in LED illuminated growth cabinets at 16°C and 26°C. Under these conditions, the most striking response was found for the stem cell population as observed by *CLV3* reporter activity. Whereas in plants grown at 26°C the strongest *CLV3* reporter signal could be observed in the stem cells of the L1 layer (Figure 4: A,B), the highest signal intensity of the reporter at 16°C was consistently found in the stem cells of the L2 layer indicating an apical shift of *CLV3* activity upon exposure to higher ambient temperature (Figure4: A-D). To analyze this phenomenon more quantitatively for a group of individual plants, we used the nuclear segmentation to calculate the mean pixel intensity for the *CLV3* reporter signal in every nucleus in the shoot meristem. Only nuclei inside spherical sections were analyzed (Figure 4: E,F). This allowed us to quantify reporter activity on a single cell level. However, since fluorophore maturation depends significantly on temperature (Macdonald et al., 2012) (Balleza et al., 2018), an absolute comparison of reporter signal intensities was not possible. Therefore, for a given reporter, we chose to compare mean pixel intensities in nuclei by using the maximum value of these intensity values as an internal reference. This allowed us to compare the spatial distributions of reporter expression at different temperatures in different clonal layers of the SAM relative to the location of the respective signal intensity maximum, thereby normalizing for the effect of temperature on fluorophore maturation. In addition, we counted the number of *CLV3* positive cells for a given meristem sample by using an arbitrary threshold for the mean intensity. Here, nuclei with a mean pixel intensity of e.g. at least 35% of the maximum mean pixel intensity for the *CLV3* reporter were counted as positive and the ratio of stem cell numbers in L1 and L2 in SAMs at different temperatures were compared (Figure 4J). These ratios were below 1 (on average 0.77) indicating higher *CLV3* activity in L2 at 16°C compared to a ratio of around 2.5 for plants grown at 26°C demonstrating an obvious shift in *CLV3* activity at higher ambient temperatures within the SAM stem cell domain. To investigate the overall SAM morphology of plants grown at different temperatures we analyzed SAM size and cellular architecture of spherical sections as described. The radius of spherical SAMs decreased by more than 30% in plants cultivated at 26°C (Figure4, G). At the same time the number of L1 cells in spherical sections with the same outer surface size increased by about 20% whereas the number of L2 cells was reduced by almost 30% (Figure4, H) resulting in a consistent change of the respective cell number ratios from around 1.2 to 2 (Figure4, I). This meant that at 16°C the central region of the SAM had almost the same cell density in L1 and L2 whereas at 26°C there were twice as many L1 cells compared to the L2 layer. Thus, the apical shift of *CLV3* reporter activity correlated with a layer specific change in cell numbers, which could be indicative for a corresponding layer dependent change in cell proliferation rates.

**Figure 4:**
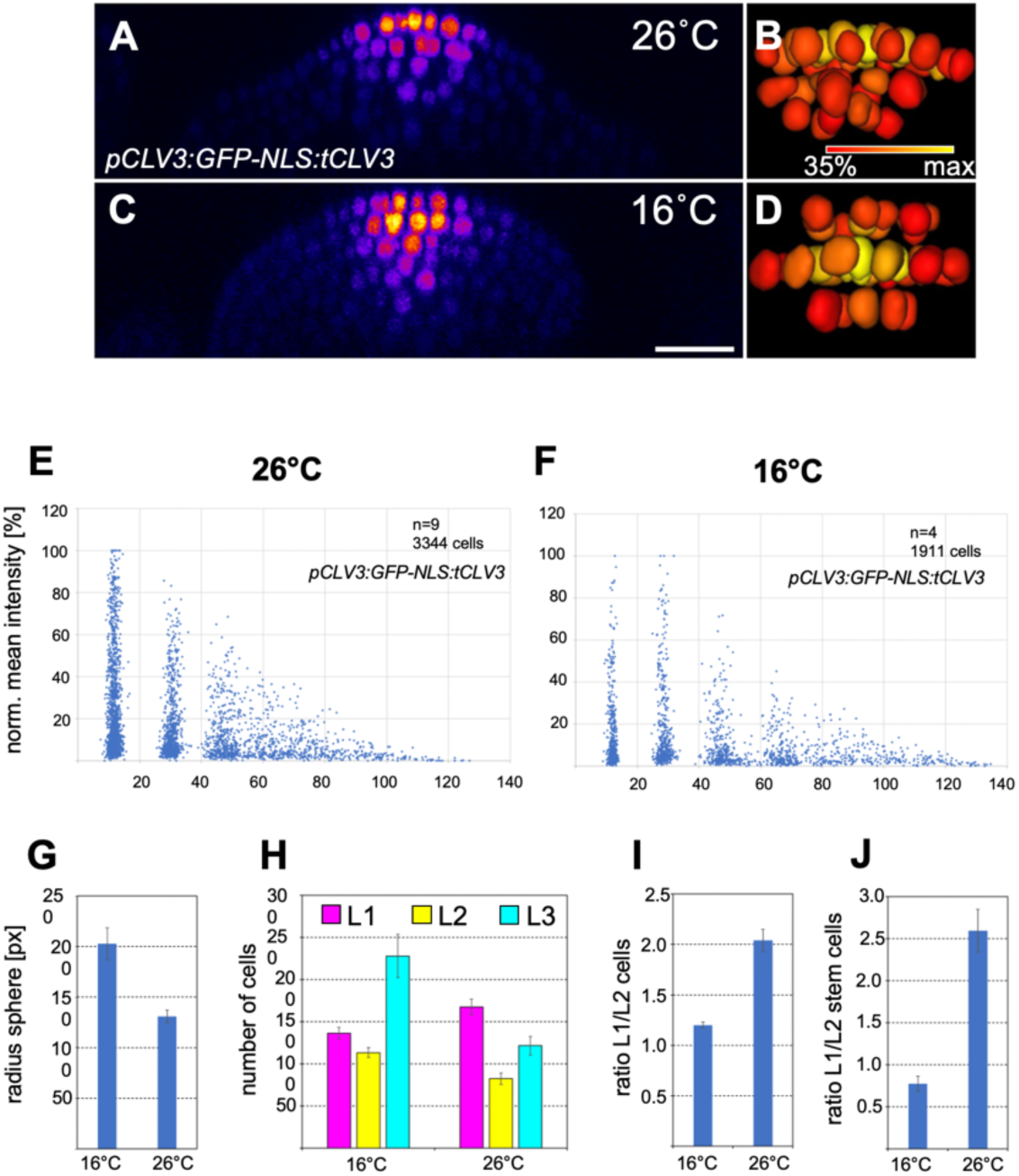
Ambient temperature affects spatial distribution of *CLV3* reporter signal. Double reporter plants (*pUBQ10:3xmCherry-NLS:tUBQ10* and *pCLV3:GFP-NLS:tCLV3*) grown continuously at 16 and 26°C in LD conditions were imaged and analyzed. Cross sections of representative meristem samples are shown in A (26°C) and C (16°C) using the Fiji Fire lookup table. At 26°C the maximum of *CLV3* reporter expression can be observed in the stem cells of the L1 layer whereas at 16°C the reporter activity is highest in L2 stem cells. B and D show 3D reconstructions of respective stem cell domains from such meristem samples with heatmaps of *CLV3* reporter signal. Only nuclei that had at least 35% of the highest mean pixel intensity value are displayed. Scalebar in A-D (length of heatmap legend in B and D) corresponds to 20 µm. To monitor *CLV3* reporter activity for a group of individual plants cultivated at different ambient temperatures the mean pixel intensity was calculated for each nucleus within a spherical section. Values were min-max normalized for each meristem and are collectively plotted against the distance of each individual nucleus to the outer meristem surface for 26°C (E) and 16°C (F). The shift of maximum *CLV3* reporter signal from L1 to L2 can be observed. G: Radius of the spherical SAM is displayed for plants cultivated at 16°C and 26°C. The number of cells per layer in spheric sections with equal outer surface size is shown in H for two different temperatures. Respective ratios of cell numbers in L1 and L2 i was calculated and is shown in I. Based on mean GFP pixel intensities per nucleus stem cells were counted as described. Ratio of stem cells in L1 and L2 is shown in J for plants grown at 16°C and 26°C.

### Molecular patterning of the stem cell domain is fixed around floral transition

To analyze the effect of changes in ambient temperature on the dynamics of reporter activity patterns we cultivated triple reporter plants at 26°C for various periods of time before plants were shifted to 16°C. Inflorescence meristems of such plants were imaged and analyzed using our newly developed pipeline. In control plants continuously grown at 26°C the strongest *CLV3* reporter signal was again observed in the epidermal stem cell layer. Interestingly, under these conditions, we could also observe a number of cells in the L2 layer with weak but significant *WUS* reporter signal (Figure5: A’,A”). The pattern for the *CLV3* reporter was similar in plants kept at 16°C for 10 days after initial 20 days at 26°C and for plants kept at 16°C for 19 days after initial exposure to 26°C for 21 days (Figure 5: B’,C’). However, the *WUS* reporter signal observed in L2 cells at higher ambient temperature was gradually reduced in response to cold temperature exposure (Figure 5: B”,C”). In contrast to the stability of *CLV3* reporter activity in plants that experienced at least 20 days of 26°C, we found a dynamic response of the stem cell domain in plants that had been exposed to warm temperatures for a shorter amount of time. The maximum reporter activity could be predominantly observed in L2 cells in plants grown at 26°C for 13 days followed by 25 days exposure at 16°C (Figure5: D’) and it had completely shifted to the L2 layer after cultivating for 10 days at 26°C followed by 32 days at 16°C (Figure5: E’). In both experiments no significant *WUS* reporter signal could be observed in L1 or L2 cells (Figure5: D”,E”). Since under these conditions, *Arabidopsis* plants undergo the transition from vegetative growth to flowering around 10-14 days, these findings suggested that the molecular patterning of the stem cell domain was fixed prior to, or during floral transition.

**Figure 5:Temperature.**
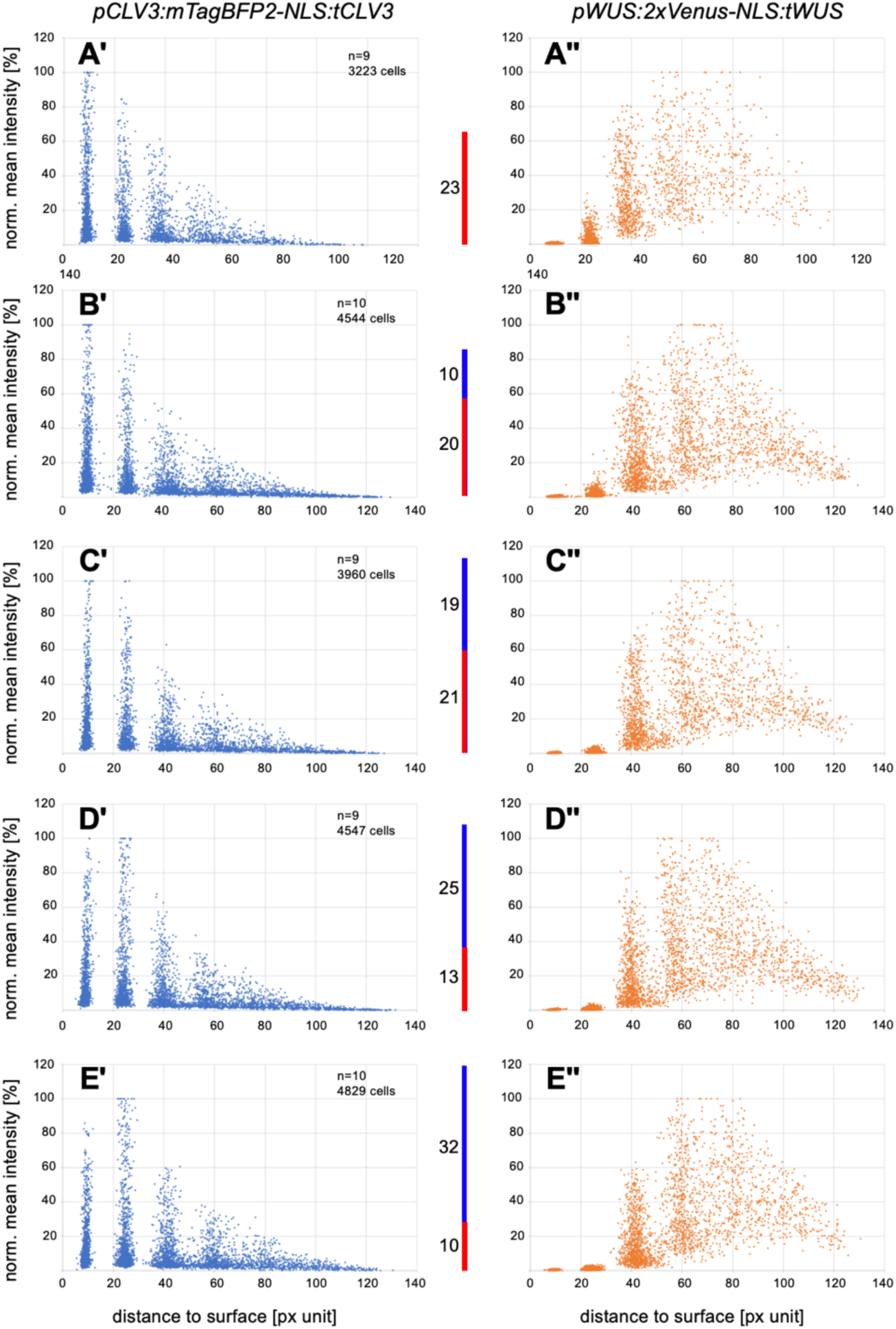
shift experiments reveal dynamic responses of *WUS* and *CLV3*. Triple reporter plants (*UBQ10, CLV3, WUS*) were cultivated for different time periods at 26°C before they were shifted to 16°C (number of days at 26°C (red) and at 16°C (blue) as indicated in vertical bars). SAMs were imaged after inflorescences had reached a shoot height of about 10-20 cm. Mean signal intensity for *CLV3* reporter (A’-E’) and *WUS* reporter (A”-E”) were measured for all nuclei in central spherical sections for the indicated number of plants. A’,A”: 23 days at 26°C (no shift), B’,B”: 20d/10d, C’,C”: 21d/19d, D’,D”: 13d/25d, E’, E””: 10d/32d. If plants were exposed to warm ambient temperature until bolting started, the maximum of *CLV3* reporter signal could be consistently observed in L1 cells even after prolonged cultivation at 16°C (A’-C’). The situation was different if plants had been shifted earlier in development. Here the reporter showed highest mean pixel intensity in nuclei located in L2. These changes in reporter activity were obvious if the temperature shift occurred before or around the flower transition (D’,E’). The *WUS* reporter shows no signal in L1 and a weak signal in L2 nuclei in plants kept at 26°C without temperature shift (A”). The L2 signal gradually disappears if plants were shifted to 16°C and kept for increasing time periods.

### Ambient temperature affects relative cell numbers in meristem layers

To address whether ambient temperature also had an effect on the overall morphology and cellular architecture of the SAM, which may correlate with changes in the stem cell domain, we compared the overall size of the meristem as well as the number of cells in different clonal layers in spherical sections after time shift experiments (Figure6: A,B,C). Whereas plants continuously grown at 26°C had significantly smaller meristems (mean radius = 134 px) the radius of the SAM was larger for all plants that have been shifted for different periods to 16°C (ranging from 172 to 191 px, Figure6: A). This result showed, that while the morphology of the stem cell domain is fixed at a certain developmental time, SAM size is not, allowing the meristem to respond to variations in ambient temperature throughout the life cycle. This suggested that stem cell fate and SAM activity are uncoupled by environmental inputs.

To determine how this may be achieved at the cellular level, we next analyzed the cellular architecture of our specimen by using spherical sections containing stem cells and part of the surrounding transient amplifying cells (see Figure S1). Using this approach, we observed that the number of L1 cells showed only minor variations. There were on average between 143 and 153 L1 cells in all groups of investigated plants. This result was slightly different then what we had observed in plants grown without temperature shift where we could observe a 20% increase in L1 cell number at 26°C compared to 16°C. The observed differences in SAM size were thus largely accompanied by changes in L1 cell size. Interestingly, the number of L2 cells gradually increased on average from 73 to 111 depending on the timepoint of temperature shift. Accordingly, the ratio of L1 to L2 cells gradually decreased from 2 in plants grown at 26°C to 1.29 in plants that had been grown for 10 days at 26°C followed by 32 days at 16°C (Figure 6: C). These results were very similar to what we could observe under static temperature conditions. Hence in our shift experiments, in contrast to the epidermis, temperature responses of the L2 layer could involve changes in cell proliferation. Consistently, we found similar changes when specifically analyzing stem cells: When we determined the ratio of stem cells in the L1 and L2 layer (Figure 6: D) we found the ratio between L1/L2 cell numbers to decrease from 2.59 to 0.75 reflecting again the observed apical shift of *CLV3* reporter expression at 26°C (Figure6: E).

**Figure 6:**
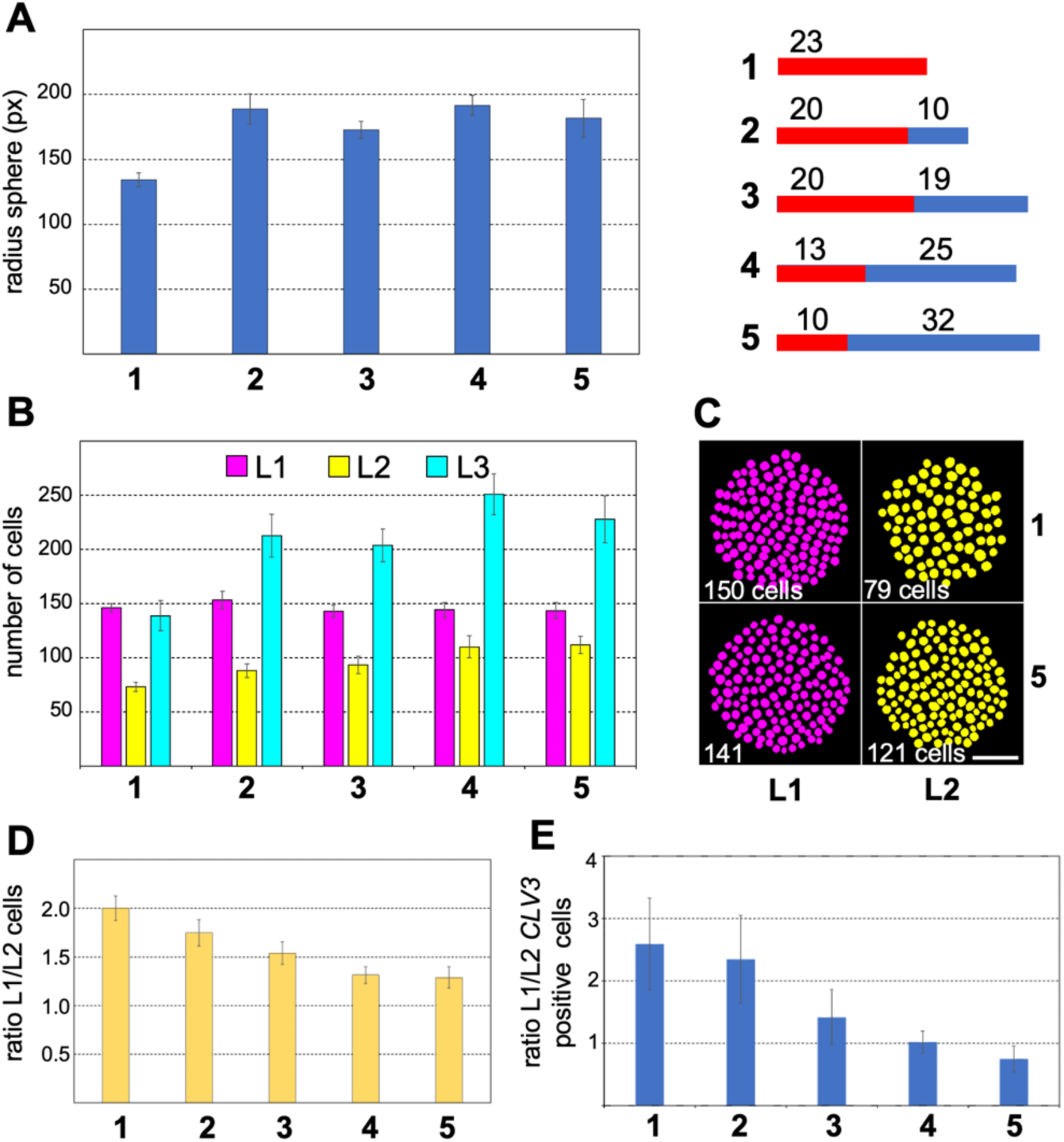
Changes in ambient temperature affect tissue layer specific cell proliferation. The data from our temperature shift experiments were used to analyze dynamic morphological changes in the SAM during temperature adaptation. The size of meristems is given in (A) by the radius of the fitted sphere. The meristems of plants kept at 26°C were significantly smaller than those that had been shifted to 16°C. The number of cells per layer in spherical sections of equal surface size (excluding the cone surface) are displayed in B. Whereas the number of L1 remains relatively constant the number of L2 cells gradually increases. The corresponding decreasing L1/L2 ratios of cell numbers is depicted in D. A representative example of how cell densities are affected by ambient temperature is shown in C. Here, max projections of L1 and L2 nuclei of equally sized sections from 2 individuals of groups 1 and 5 are shown. Scalebar corresponds to 20 µm. E: The numbers of stem cells in different layers were counted based on *CLV3* reporter signal intensities as described and the L1/L2 ratios of stem cell numbers is shown. Ratios below 1 are indicative for the observed L1 -> L2 shift in maximum *CLV3* reporter activity.

### The spatial pattern of WUS-GFP fusion protein is only slightly affected by changes in ambient temperature

It has been reported that *CLV3* expression is directly controlled by *WUS* (Perales et al., 2016). Since WUS protein moves from the cells of the OC to the stem cells via plasmodesmata (Daum et al., 2014; Yadav et al., 2011), we asked if the relative WUS distribution within the meristem might be affected by different temperatures (Figure7). This could potentially explain the observed temperature dependent change in *CLV3* reporter activity. We therefore analyzed our well established WUS-GFP rescue line in which endogenous WUS has been replaced by a *pWUS:GFP-linker-WUS:tWUS* transgene (Daum et al., 2014). We found minor differences in WUS-GFP distribution in plants grown constantly at 26°C or 16°C using the spatial distribution of cells based on normalized mean pixel intensity values for all nuclei in spherical section (Figure7: A,B).

To measure the overall distribution of WUS-GFP signal in the inflorescence SAM we used the sum of all pixel intensity values for every nucleus in a cylinder with equal radius that surrounds the central axis of the SAM and monitored the relative distribution of the signal in the different clonal layers of the meristem (Figure 7: C). The relative proportion of WUS-GFP in the L1 and L2 layers seemed to be slightly increased at 26°C (11.8% vs 7.2% in L1 cells and 18.5% vs 14.3% in L2 cells). This small difference could be a consequence of the minimal apical shift in *WUS* promoter activity observed in our temperature shift experiments (Figure5: A”). In sum, the temperature dependent variations in the spatial expression pattern of the stem cell specific *CLV3* gene as defined by *CLV3* reporter activity were much more pronounced than the changes in distribution of WUS-GFP protein. Thus, it remained open if temperature dependent changes in WUS protein mobility could be the cause for the observed changes in *CLV3* reporter activity, or if other mechanisms were at play.

**Figure 7:**
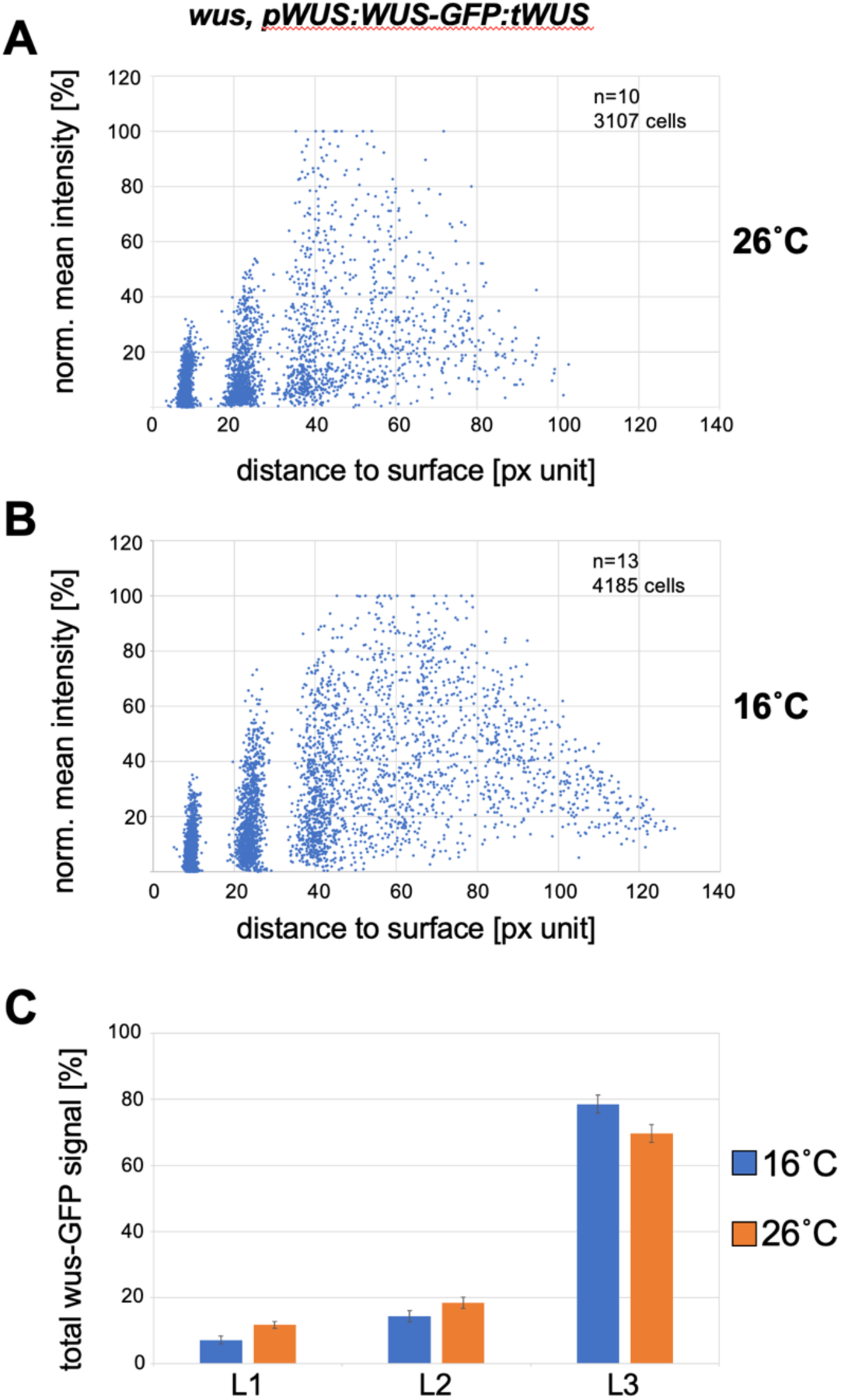
Layer specific levels of WUS-GFP are only moderately affected by temperature. To investigate if ambient temperature influences the cell to cell mobility of the WUS transcription factor we used a *wus* mutant line expressing WUS-GFP under the *WUS* promoter (Daum et al., 2014). Plants were grown at 26°C or 16°C before meristems were harvested and imaged. Spherical sections of groups of plants were analyzed and the results are shown in A and B. Since the *CLV3* reporter was not present in this line we used the WUS-GFP signal in L2 nuclei to determine the summit point of the meristems. To quantify the relative proportion of WUS-GFP in different cell layers we used the sum of pixel intensities for each nucleus within a cylinder of radius r=70 px around the central axis of the meristem. Using this approach, we obtained relative distributions of WUS-GFP for all three layers as shown in C.

## Discussion

The plant shoot apical meristem is a developmental motor underlying postembryonic growth by fueling organ formation. At the same time it integrates information to synchronize its activity with environmental conditions and availability of resources. To understand the nature and dynamics of such processes, cell behavior and gene expression must be precisely monitored under diverse growth conditions.

Here we presented an imaging-based approach to study acclimatization processes in the shoot apical meristem. We used different genetically encoded reporter constructs that allow extraction of general morphological parameters as well as the quantification of specific reporter gene activities to monitor the activity of the stem cell system in the SAM.

In contrast to many studies that analyzed plant cell morphologies in complex tissues (Federici et al., 2012; Kierzkowski et al., 2019; Montenegro-Johnson et al., 2019, 2015; Wolny et al., 2020) we used nuclear targeted fluorophore based reporter constructs instead of cell wall staining. Despite being more time consuming in generating and analyzing such reporter lines, we found that this approach had several advantages. Firstly, we experienced difficulties in detecting cells in deeper SAM layers when using cell wall staining dyes or genetically encoded cell wall markers. In contrast, nuclear signals were much brighter and more robust allowing a reliable imaging of higher numbers of respective samples in a highly reproducible way. Secondly, the analysis of images of fluorophores targeted to the nucleus was straight forward and much more robust than identifying cell outlines. Due to advances in AI assisted 3D segmentation algorithms a major step of data processing could be carried out with high precision without much human input. It allowed the extraction of SAM size as well as total cell numbers and size of subpopulations of cells within the meristem. In the small, largely undifferentiated cells of the shoot apical meristem nuclear size can also be used as a good approximation for cell size (see Figure S2). We could show that nuclear and cell volumes show high linear correlation which allows to indirectly deduce information about cell sizes by just analyzing sizes of the nuclei.

Our experiments revealed subtle changes in the spatial expression pattern of *CLV3* and *WUS* reporters when plants were cultivated under different ambient temperatures. Expression of both reporters was shifted apically in warm temperatures. The effect was stronger for the *CLV3* reporter whose expression maximum shifted from L2 stem cells to L1. For plants that had been initially cultivated at high temperatures and were later shifted to 16°C the apical shift of *CLV3* expression was most prominent if the temperature shift occurred before or during floral transition, suggesting that the window of developmental plasticity closes around this time. In line with the apical shift of *CLV3* promoter activity, the *WUS* reporter showed a weak signal in L2 cells in SAMs cultivated only at 26°C which could not be detected in plants that had been exposed to 16°C for longer time periods.

Since CLV3 is a small secreted peptide that acts non cell-autonomously it seemed unlikely that the shift of maximum *CLV3* expression from L1 to L2 has major direct consequences for stem cell homeostasis. It has been proposed that the dosage of *CLV3* expression can vary over one order of magnitude without visible phenotypical consequences for meristem development (Müller et al., 2006).

We found a correlation of the apical shift of stem cell reporter signals with the layer specific cellular architecture of the SAM in response to increased temperature. The ratio of L2 to L1 cells in the central section of the SAM gradually decreases after plants are shifted from 26°C to 16°C. The number of L1 cells remains relatively constant whereas the number of L2 cells gradually increases by more than 30%. This increase in L2 cell number could point to a mechanism that controls cell proliferation in the central domain of the SAM in a layer specific manner. Alternatively, the observed changes could be explained by a global shift towards differentiation meaning that the balance between formation of new cells in the central zone and consumption of cells in the peripheral zone would be affected. Such processes can be observed during SAM senescence where stem cells gradually lose their ability to divide while new organs are still formed at the periphery. This leads to a gradual decrease in SAM size and eventually to SAM arrest (reviewed in (Wang et al., 2023)). It seems unlikely however that such a scenario could lead to the layer specific changes in cell numbers in the central part of the SAM as it is observed in our study. It would be interesting to apply our analysis to changes in SAM architecture observed during aging. From our imaging data we have no evidence for enhanced rates of cell death which could also explain the local differences in nuclear densities in response to temperature changes which leaves the layer specific local adaptation of cell proliferation rates as the most likely explanation.

Overall, it remains unclear if this change in cellular tissue morphology has a physiological function. It has been reported that genetically induced delay of cell proliferation in the SAM stem cell domain can be corrected by compensatory mechanisms in the periphery of the SAM. It was also shown that such perturbations can however affect the emergence of organ primordia (Serrano-Mislata et al., 2015). We do observe a higher rate of organ formation in plants grown at colder temperatures (data not shown) but it remains unclear if this can be attributed to the very specific changes in cellular architecture of the central meristem.

Nevertheless it would be interesting to see if initially reduced numbers of L2 cells in the central part of the meristem could be compensated by higher L2 proliferation rates at the periphery. WUS was shown to be a mobile transcription factor that moves via plasmodesmata from the cells of the OC to the stem cells where it induces stem cell fate and as a result the expression of *CLV3* (Daum et al., 2014; Yadav et al., 2011). It has been reported that PDs in maize are affected by cold temperature (Bilska and Sowiński, 2010). Therefore, we investigated if the relative spatial distribution of a WUS-GFP fusion protein in the SAM is affected by ambient temperature. We could detect a small but significant change in the relative accumulation of WUS-GFP in the L1 and L2 layers of respective plants grown at higher temperatures. However, this difference was much smaller than the one observed for *CLV3* promoter activity, suggesting that additional mechanisms were at play, or that the response of the *CLV3* regulatory sequences to WUS is non-linear.

The spatial expression of *CLV3* has been shown to also depend on an apical-basal gradient of genes of the *HAIRY MERISTEM* family (Zhou et al., 2018, 2015). Epidermally expressed HAM1-GFP led to a strongly reduced *CLV3* reporter signal in the L1 and to enlarged meristems proving that there might be a functional significance for high levels of *CLV3* expression in epidermal SAM cells. Enlarged meristems usually produce more organs. Temperature dependent spatial fine tuning of *CLV3* expression in the L1 could thus be a way to adapt meristem size and as a consequence organ formation capability in response to ambient temperature changes.

The molecular mechanisms that govern the observed changes in SAM architecture described here remain to be investigated. We have analyzed *CLV3* reporter activity in *PIF4* as well as in *ARP6* mutant plants at different ambient temperatures and observed similar transcriptional and morphological responses as in wildtype (data not shown). Thus, the temperature adaptation processes in the inflorescence meristem seem to be controlled by different mechanisms.

## Materials and Methods

### Plant material

All plants were cultivated on soil at 16°C or 26°C in Polyklima S2LED6 incubators under long day conditions.

### Transgenic lines

We established a double reporter line harboring a *UBQ10* and a *CLV3* reporter in the *Col-0* background. This line was transformed with pCW119. The triple reporter line was created by transforming pCW098 (*pCLV3:mTagBFP2-NLS:tCLV3*) into a *Col-0* background harboring stable single insertions for pCW039 (*pWUS:2xVenus-NLS:tWUS*) and pCW066 (*pUBQ10:3xmCherry-NLS:tUBQ10*). The WUS rescue line wus; *pWUS::WUS-linker-GFP:tWUS* was described in (Daum et al., 2014).

### Cloning

Four different transformation vectors were cloned using green gate assembly. Details about the cloning procedure can be found in the supplement. Modules marked with * have been described in (Lampropoulos et al., 2013) modules marked with ** were described in (Schürholz et al., 2018). For some experiments we used pCW119, a construct that combined the *UBQ10* reporter with a *CLV3* reporter on the same T-DNA.

### Microscopy

Inflorescence tips were collected at a shoot height of around 10-20 cm and mounted in 2% Agarose containing petri dishes. Obstructing flower primordia were removed so that the meristem was exposed and the plate was subsequently filled with water so that all samples were completely submerged. For cell segmentations meristems were stained with DAPI (200µg/ml) for 3 minutes prior to imaging. Image stacks were recorded using an upright Nikon A1 Confocal with a CFI Apo LWD 25 × water immersion objective (Nikon Instruments). Pools of approximately 10 plants were imaged for each experiment with the same settings. Image stacks were always recorded with the same dimension (164,41µm x 164,41µm x 45µm). Accordingly voxel size of recorded images was 0.32×0.32×0.5µm. For analysis image stacks were made isotropic (0.32×0.32×0.32µm) in Fiji using bicubic interpolation.

### Image analysis

Analysis of 3D image volumes was performed in KNIME Analytics platform 4.5.0 with following extensions: KNIME Image processing extension, KNIME Matlab scripting extension, KNIME Python scripting extension, KNIME ilastik integration and KNIME ImageJ integration. To run the workflows a local installation of Matlab (we used R2018b), Python (3.9) and ilastik (1.4.0rc6) is required. In addition, the ejml java library (v.0.23) was used for the sphere fitting part and had to be added as external library. KNIME workflows were run on a Linux Workstation with Ubuntu 22.04.2 LTS and on MacBook Pro with macOS 11.5.2.

Nuclear segmentation of the *pUBQ10:3xmCherry-NLS:tUBQ10* channel was obtained using StarDist (“StarDist - Object Detection with Star-convex Shapes,” 2023). For training a respective model we had to generate a ground truth. Here we used 8 randomly generated substacks (160×160×64) from each of 4 different *pUBQ10:3xmCherry-NLS:tUBQ10* isotropic image stacks (512×512×142). For each set of these smaller image stacks we obtained a corresponding nuclear segmentation using the seeded watershed function in KNIME with manual seeding. We used 8 pairs of small image-segmentation stacks to train our model on Google Colabs Server using the respective StarDist example jupyter notebook for 3D segmentation (https://github.com/stardist/stardist/blob/master/examples/3D/2_training.ipynb). This solution was chosen because of the computational demands of the training process. We found that using image-segmentation pairs from the same larger image stack gave better segmentation results then using mixed sets. The model predictions to obtain the final full size nuclear segmentations of all analyzed image stacks were run locally in KNIME using the Python scripting integration and modified code from the respective StarDist example jupyter notebook (https://github.com/stardist/stardist/blob/master/examples/3D/3_prediction.ipynb). To generate 3D binary masks from the *pUBQ10:3xmCherry-NLS:tUBQ10* channel we trained a pixel classification model in ilastik (using all features with sigma 5 and 10). Using just one single stack to train the pixel classification model was usually sufficient. However, it was sometimes necessary to train a new model using a different image volume to accommodate for changes in image acquisition or sample type. The predictions for the binary masks for all image stacks were generated along with the StarDist predictions in a designated KNIME workflow using the ilastik integration and the respective ilp project file. The rest of the analysis was done in the main KNIME analysis workflow. This workflow allows to observe and correct certain steps of the analysis interactively. It also saves results as files containing quantitative data as well as special image stacks for visual inspection.

The summit point of the meristem (with coordinates x_s_ y_s_ z_s_) was determined by averaging the coordinates of the centroids of the 20 L1 nuclei (L2 in case of the WUS-GFP line) with the highest mean pixel intensity for the clv3 (WUS-GFP) channel:

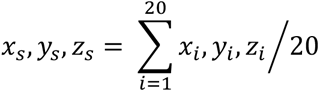

The apico-basal axis of the meristem was defined as the line passing through the center point of a sphere fitted through the centroids of L1 nuclei at a user defined distance D_s_ from the summit point. D_s_ was kept constant (100 px) for most of the experiments and was only adapted for very small meristems (90 px).

Visualization of 3D image volumes was done using Fiji 3D viewer. The cell segmentation for Figure S2 was done in KNIME using 3D seeded watershed with a boundary prediction image created with PlantSeg (Wolny et al., 2020) and our nuclear segmentation as seeds. Two annotated KNIME workflows (ilastik_mask.knwf and Triple Reporter Workflow.knar.knwf and an ImageJ macro that facilitates the manual drawing of Bezier lines to separate primordia from meristem were deposited on Zenodo (Zenodo_Repo DOI: 10.5281/zenodo.7892435).

## Funding

This work was supported by the DFG through funds to the SFB873 project B01.

## Supplemental Material

### Cloning of transcriptional reporter constructs

Four different transformation vectors were cloned using green gate assembly. Modules marked with * have been described in (Lampropoulos et al., 2013) modules marked with ** were described in (Schürholz et al., 2018). For all remaining module-vectors the sequences of oligos used for PCR amplification of the respective module are given below:

**- pCW039 (*pWUS:2xVenus-NLS:tWUS*):**

(pGG-A003*, pCW018, pCW019, pCW016, pGG-E002*, pGG-F003* -> pGG-Z002) pCW018:

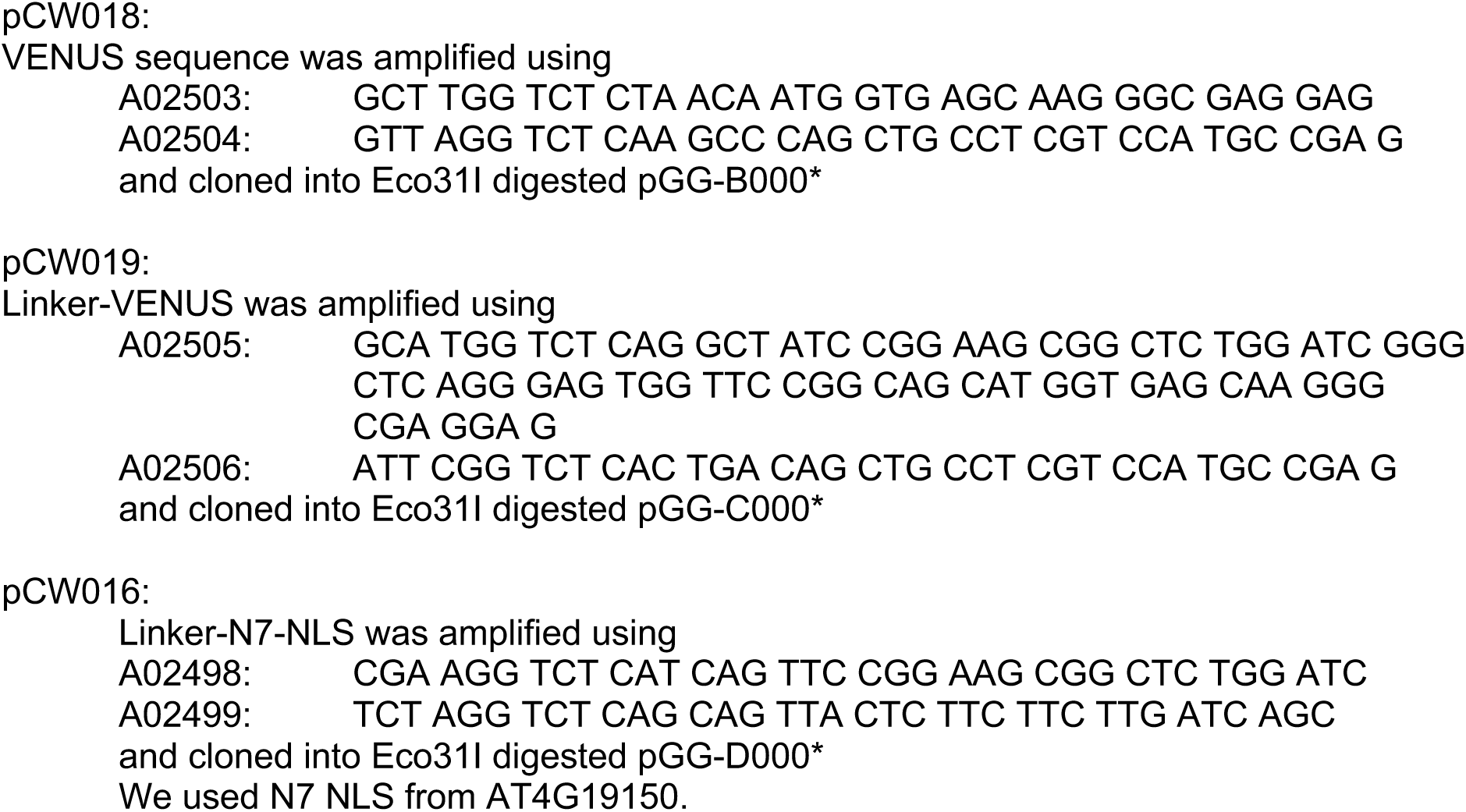

pGG-Z002 is a derivative of pGG-Z003* where LB and RB sequences were swapped.

**- pCW098 (*pCLV3:mTagBFP2-NLS:tCLV3*):**

(pGG-A033**, pGG-B002*, pCW091, pGG-D007*, pGG-E008**, pGG-F001* -> pGG-Z003*): pCW091 (mTagBFP2):

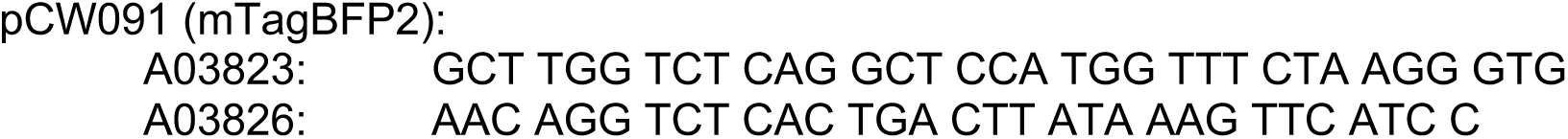

**- pCW066 (*pUBQ10:3xmCherry-NLS:tUBQ10*):**

(pGG-A006*, pGG-B003*, pGG-C026*, pGG-D007*, pGG-E009*, pGG-F007* -> pGG- Z003*)

**- pCW119** (***pUBQ10:3xmCherry-NLS:tUBQ10, pCLV3:GFP-NLS:tCLV3):***

(pGG-A033**, pGG-B003*, pGG-C014*, pGG-D007*, pGG-E008**, pGG-F001* -> pCW086)

pCW086:

This is a derivative of pGG-Z003. pUBQ10:3xmCherry-NLS:tUBQ10 was amplified using oligos A04166 and A04167 and pCW066 as template. The PCR product and pGG-Z003 were digested with AscI and ligated.

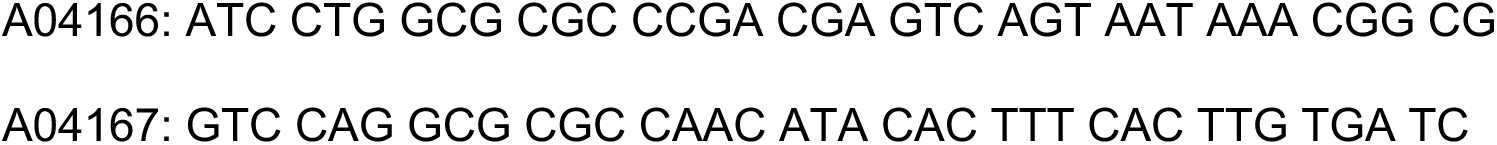

**Figure S1.**
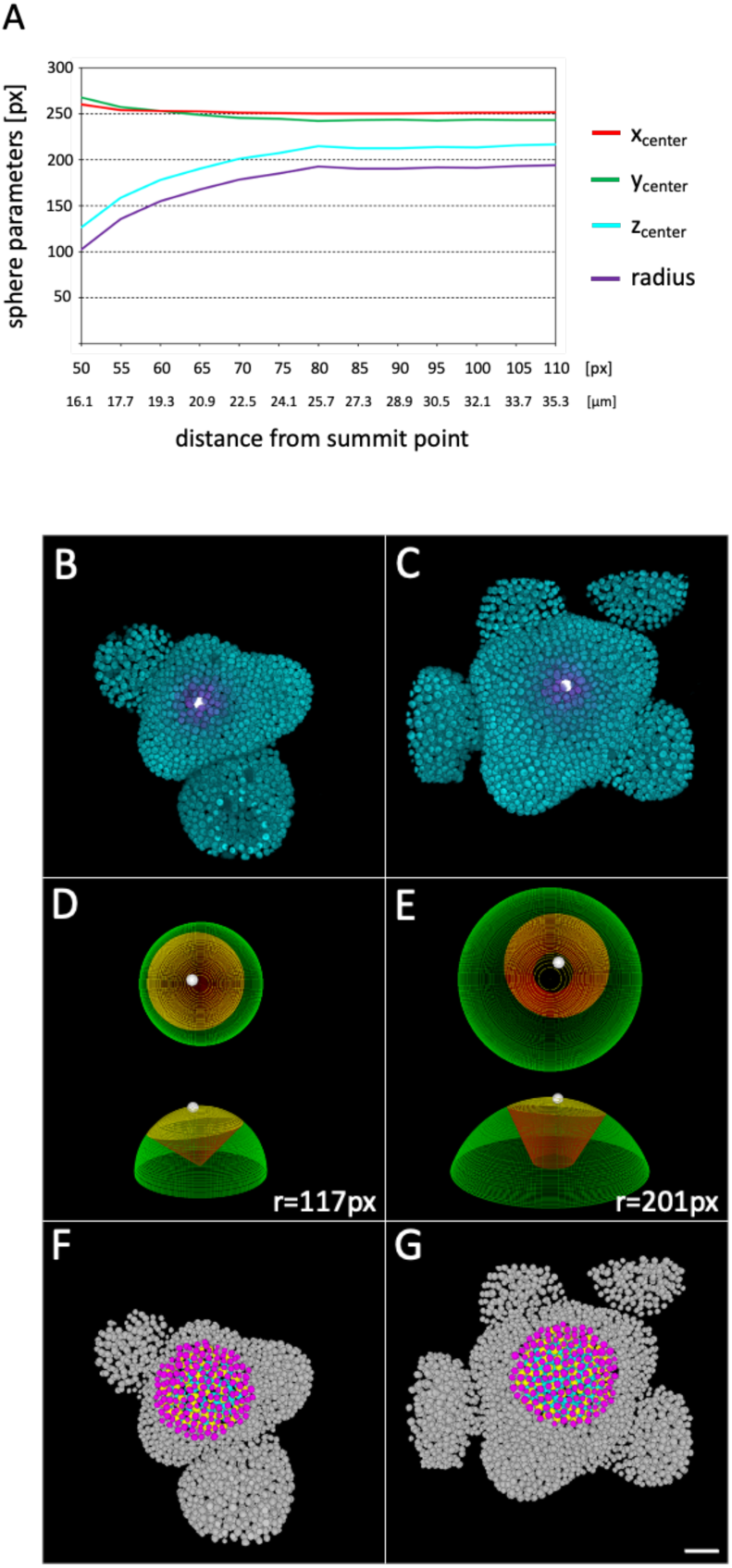
A: Effect of distance from summit point on sphere fitting parameters. Increasing the distance would lead to more points (or centroids of L1 nuclei) used as input for the sphere fit. At small distances the fitting algorithm suggests spheres that are too small whereas at distances greater than 80px (=25µm) the coordinates of the center point of the sphere as well as the sphere radius converge stably and follow the size of the dome shaped meristem very closely. We used distances of 90 or 100 px units for all experiments. Data from one representative meristem are shown. B,C: 3D reconstructions of the smallest and the largest meristem in our triple reporter dataset with *pCLV3:mTagBFP2-NLS:tCLV3* (magenta) and *pUBQ10:3xmCherry-NLS:tUBQ10* (cyan). The *CLV3* signal was used to define the summit point of the meristem (small white sphere, see materials and methods section). D,E: top and side views of respective fitted spheres (green). The corresponding radius is displayed in px units. Spheric sections (red) were selected for both spheres with equally sized spherically curved surface area (yellow region). To this end the user can define the size of this surface area (here it was 30000 px^2^ which corresponds to 3093 µm^2^) and the workflow will automatically calculate the corresponding elevation angle threshold (see Figure 3: D2,D3) required for selecting nuclei for downstream analysis. F,G: Resulting selections are shown in color (L1: magenta, L2: yellow, L3: cyan) for both meristems. Gray nuclei were excluded from the reporter analysis. Scalebar corresponds to 20 µm.

**Figure S2.**
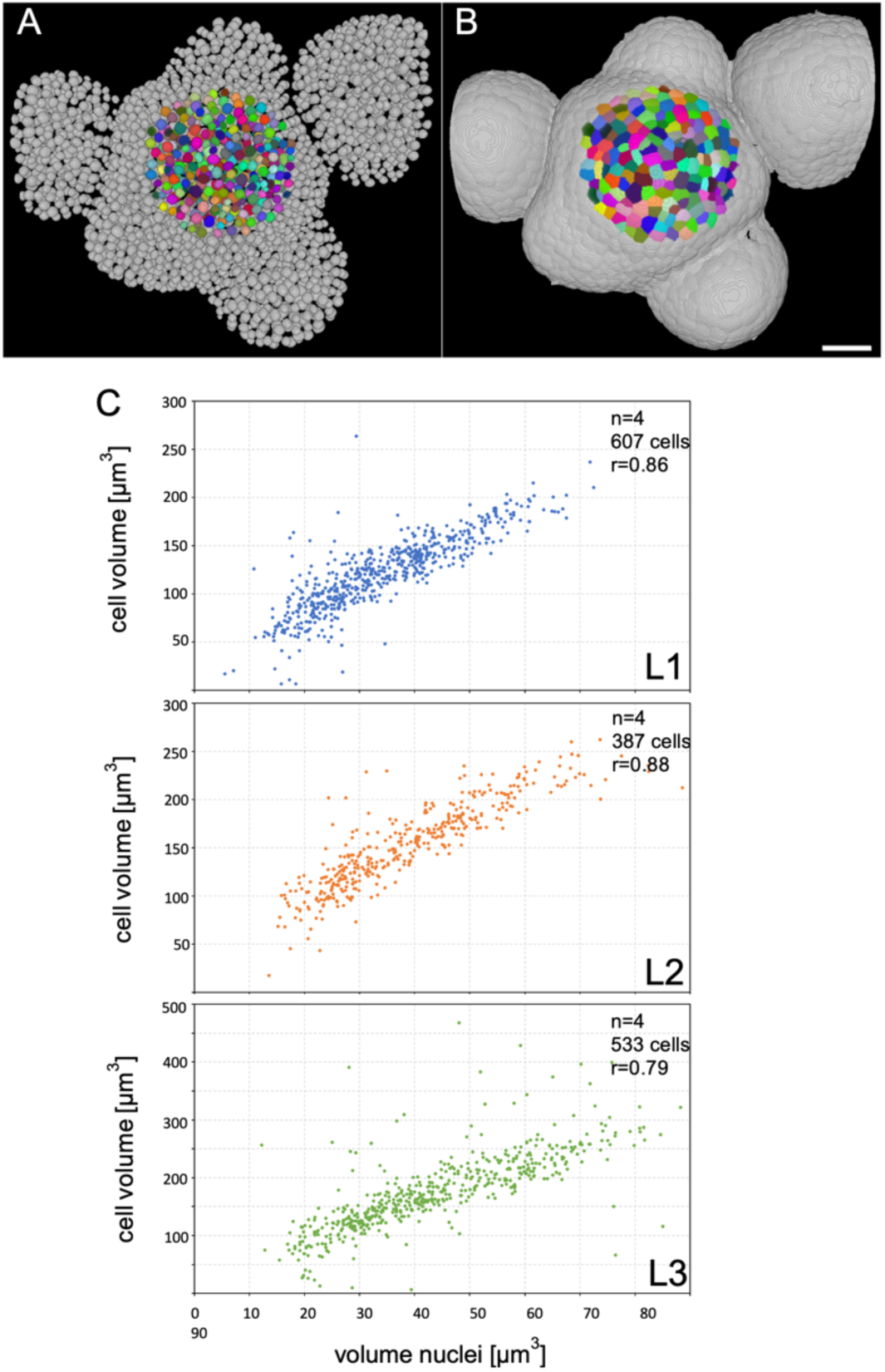
Linear Correlation between nuclear size and cell size in the SAM Inflorescence meristems expressing *pUBQ10:3xmCherry-NLS:tUBQ10* were prepared for imaging and treated with DAPI to stain the cell wall for obtaining cell segmentations. A: Nuclear segmentation. Spherical sections (colored nuclei) were used to select nuclei for volume analysis (gray nuclei were not included in this analysis). B: corresponding cell segmentation. Scalebar corresponds to 20 µm. C-E: Nuclear and cell volumes were plotted for L1, L2 and L3 cells of 4 different meristems grown under long day conditions at 21°C. Numbers of analyzed cells and Pearson correlation coefficients (r) are displayed. Many of observed outliers can be attributed to segmentation errors which are more frequent in L3 cells due to the decreasing signal to noise ratio in deeper cell layers.

## Notes

### Competing Interest Statement

The authors have declared no competing interest.

## References

Balleza, E., Kim, J.M., Cluzel, P., 2018. Systematic characterization of maturation time of fluorescent proteins in living cells. Nat. Methods 15, 47–51. https://doi.org/10.1038/nmeth.4509

Bartrina, I., Otto, E., Strnad, M., Werner, T., Schmülling, T., 2011. Cytokinin regulates the activity of reproductive meristems, flower organ size, ovule formation, and thus seed yield in Arabidopsis thaliana. Plant Cell 23, 69–80. https://doi.org/10.1105/tpc.110.079079

Berg, S., Kutra, D., Kroeger, T., Straehle, C.N., Kausler, B.X., Haubold, C., Schiegg, M., Ales, J., Beier, T., Rudy, M., Eren, K., Cervantes, J.I., Xu, B., Beuttenmueller, F., Wolny, A., Zhang, C., Koethe, U., Hamprecht, F.A., Kreshuk, A., 2019. ilastik: interactive machine learning for (bio)image analysis. Nat. Methods 16, 1226–1232. https://doi.org/10.1038/s41592-019-0582-9

Berthold, M.R., Cebron, N., Dill, F., Gabriel, T.R., Kötter, T., Meinl, T., Ohl, P., Thiel, K., Wiswedel, B., 2009. KNIME - the Konstanz information miner: version 2.0 and beyond. ACM SIGKDD Explor. Newsl. 11, 26–31. https://doi.org/10.1145/1656274.1656280

Bilska, A., Sowiński, P., 2010. Closure of plasmodesmata in maize (Zea mays) at low temperature: a new mechanism for inhibition of photosynthesis. Ann. Bot. 106, 675– 686. https://doi.org/10.1093/aob/mcq169

Brand, U., Fletcher, J.C., Hobe, M., Meyerowitz, E.M., Simon, R., 2000. Dependence of stem cell fate in Arabidopsis on a feedback loop regulated by CLV3 activity. Science 289, 617–619. https://doi.org/10.1126/science.289.5479.617

Brand, U., Grünewald, M., Hobe, M., Simon, R., 2002. Regulation of CLV3 Expression by Two Homeobox Genes in Arabidopsis. Plant Physiol. 129, 565–575. https://doi.org/10.1104/pp.001867

Casal, J.J., Balasubramanian, S., 2019. Thermomorphogenesis. Annu. Rev. Plant Biol. 70, 321–346. https://doi.org/10.1146/annurev-arplant-050718-095919

Daum, G., Medzihradszky, A., Suzaki, T., Lohmann, J.U., 2014. A mechanistic framework for noncell autonomous stem cell induction in Arabidopsis. Proc. Natl. Acad. Sci. U. S. A. 111, 14619–14624. https://doi.org/10.1073/pnas.1406446111

Federici, F., Dupuy, L., Laplaze, L., Heisler, M., Haseloff, J., 2012. Integrated genetic and computation methods for in planta cytometry. Nat. Methods 9, 483–485. https://doi.org/10.1038/nmeth.1940

Fiorucci, A.-S., Galvão, V.C., Ince, Y.Ç., Boccaccini, A., Goyal, A., Allenbach Petrolati, L., Trevisan, M., Fankhauser, C., 2020. PHYTOCHROME INTERACTING FACTOR 7 is important for early responses to elevated temperature in Arabidopsis seedlings. New Phytol. 226, 50–58. https://doi.org/10.1111/nph.16316

Fletcher, J.C., Brand, U., Running, M.P., Simon, R., Meyerowitz, E.M., 1999. Signaling of cell fate decisions by CLAVATA3 in Arabidopsis shoot meristems. Science 283, 1911–1914. https://doi.org/10.1126/science.283.5409.1911

Galvan-Ampudia, C.S., Cerutti, G., Legrand, J., Brunoud, G., Martin-Arevalillo, R., Azais, R., Bayle, V., Moussu, S., Wenzl, C., Jaillais, Y., Lohmann, J.U., Godin, C., Vernoux, T., 2020. Temporal integration of auxin information for the regulation of patterning. eLife 9, e55832. https://doi.org/10.7554/eLife.55832

Gruel, J., Landrein, B., Tarr, P., Schuster, C., Refahi, Y., Sampathkumar, A., Hamant, O., Meyerowitz, E.M., Jönsson, H., 2016. An epidermis-driven mechanism positions and scales stem cell niches in plants. Sci. Adv. 2, e1500989. https://doi.org/10.1126/sciadv.1500989

Jeong, S., Trotochaud, A.E., Clark, S.E., 1999. The Arabidopsis CLAVATA2 gene encodes a receptor-like protein required for the stability of the CLAVATA1 receptor-like kinase. Plant Cell 11, 1925–1934. https://doi.org/10.1105/tpc.11.10.1925

Jung, J.-H., Domijan, M., Klose, C., Biswas, S., Ezer, D., Gao, M., Khattak, A.K., Box, M.S., Charoensawan, V., Cortijo, S., Kumar, M., Grant, A., Locke, J.C.W., Schäfer, E., Jaeger, K.E., Wigge, P.A., 2016. Phytochromes function as thermosensors in Arabidopsis. Science 354, 886–889. https://doi.org/10.1126/science.aaf6005

Kierzkowski, D., Runions, A., Vuolo, F., Strauss, S., Lymbouridou, R., Routier-Kierzkowska, A.-L., Wilson-Sánchez, D., Jenke, H., Galinha, C., Mosca, G., Zhang, Z., Canales, C., Dello Ioio, R., Huijser, P., Smith, R.S., Tsiantis, M., 2019. A Growth-Based Framework for Leaf Shape Development and Diversity. Cell 177, 1405–1418.e17. https://doi.org/10.1016/j.cell.2019.05.011

Koini, M.A., Alvey, L., Allen, T., Tilley, C.A., Harberd, N.P., Whitelam, G.C., Franklin, K.A., 2009. High temperature-mediated adaptations in plant architecture require the bHLH transcription factor PIF4. Curr. Biol. CB 19, 408–413. https://doi.org/10.1016/j.cub.2009.01.046

Kumar, S.V., Wigge, P.A., 2010. H2A.Z-Containing Nucleosomes Mediate the Thermosensory Response in Arabidopsis. Cell 140, 136–147. https://doi.org/10.1016/j.cell.2009.11.006

Lambolez, A., Kawamura, A., Takahashi, T., Rymen, B., Iwase, A., Favero, D.S., Ikeuchi, M., Suzuki, T., Cortijo, S., Jaeger, K.E., Wigge, P.A., Sugimoto, K., 2022. Warm Temperature Promotes Shoot Regeneration in Arabidopsis thaliana. Plant Cell Physiol. 63, 618–634. https://doi.org/10.1093/pcp/pcac017

Lampropoulos, A., Sutikovic, Z., Wenzl, C., Maegele, I., Lohmann, J.U., Forner, J., 2013. GreenGate---a novel, versatile, and efficient cloning system for plant transgenesis. PloS One 8, e83043. https://doi.org/10.1371/journal.pone.0083043

Legris, M., Klose, C., Burgie, E.S., Rojas, C.C.R., Neme, M., Hiltbrunner, A., Wigge, P.A., Schäfer, E., Vierstra, R.D., Casal, J.J., 2016. Phytochrome B integrates light and temperature signals in Arabidopsis. Science 354, 897–900. https://doi.org/10.1126/science.aaf5656

Ma, Y., Miotk, A., Šutiković, Z., Ermakova, O., Wenzl, C., Medzihradszky, A., Gaillochet, C., Forner, J., Utan, G., Brackmann, K., Galván-Ampudia, C.S., Vernoux, T., Greb, T., Lohmann, J.U., 2019. WUSCHEL acts as an auxin response rheostat to maintain apical stem cells in Arabidopsis. Nat. Commun. 10, 5093. https://doi.org/10.1038/s41467-019-13074-9

Macdonald, P.J., Chen, Y., Mueller, J.D., 2012. Chromophore maturation and fluorescence fluctuation spectroscopy of fluorescent proteins in a cell-free expression system. Anal. Biochem. 421, 291–298. https://doi.org/10.1016/j.ab.2011.10.040

Mayer, K.F., Schoof, H., Haecker, A., Lenhard, M., Jürgens, G., Laux, T., 1998. Role of WUSCHEL in regulating stem cell fate in the Arabidopsis shoot meristem. Cell 95, 805–815. https://doi.org/10.1016/s0092-8674(00)81703-1

Montenegro-Johnson, T., Strauss, S., Jackson, M.D.B., Walker, L., Smith, R.S., Bassel, G.W., 2019. 3DCellAtlas Meristem: a tool for the global cellular annotation of shoot apical meristems. Plant Methods 15, 33. https://doi.org/10.1186/s13007-019-0413-0

Montenegro-Johnson, T.D., Stamm, P., Strauss, S., Topham, A.T., Tsagris, M., Wood, A.T.A., Smith, R.S., Bassel, G.W., 2015. Digital Single-Cell Analysis of Plant Organ Development Using 3DCellAtlas. Plant Cell 27, 1018–1033. https://doi.org/10.1105/tpc.15.00175

Müller, R., Bleckmann, A., Simon, R., 2008. The receptor kinase CORYNE of Arabidopsis transmits the stem cell-limiting signal CLAVATA3 independently of CLAVATA1. Plant Cell 20, 934–946. https://doi.org/10.1105/tpc.107.057547

Müller, R., Borghi, L., Kwiatkowska, D., Laufs, P., Simon, R., 2006. Dynamic and Compensatory Responses of Arabidopsis Shoot and Floral Meristems to CLV3 Signaling. Plant Cell 18, 1188–1198. https://doi.org/10.1105/tpc.105.040444

Perales, M., Rodriguez, K., Snipes, S., Yadav, R.K., Diaz-Mendoza, M., Reddy, G.V., 2016. Threshold-dependent transcriptional discrimination underlies stem cell homeostasis. Proc. Natl. Acad. Sci. U. S. A. 113, E6298–E6306. https://doi.org/10.1073/pnas.1607669113

Quint, M., Delker, C., Franklin, K.A., Wigge, P.A., Halliday, K.J., van Zanten, M., 2016. Molecular and genetic control of plant thermomorphogenesis. Nat. Plants 2, 15190. https://doi.org/10.1038/nplants.2015.190

Schmidt, U., Weigert, M., Broaddus, C., Myers, G., 2018. Cell Detection with Star-Convex Polygons, in: Frangi, A.F., Schnabel, J.A., Davatzikos, C., Alberola-López, C., Fichtinger, G. (Eds.), Medical Image Computing and Computer Assisted Intervention – MICCAI 2018, Lecture Notes in Computer Science. Springer International Publishing, Cham, pp. 265–273. https://doi.org/10.1007/978-3-030-00934-2_30

Schoof, H., Lenhard, M., Haecker, A., Mayer, K.F., Jürgens, G., Laux, T., 2000. The stem cell population of Arabidopsis shoot meristems in maintained by a regulatory loop between the CLAVATA and WUSCHEL genes. Cell 100, 635–644. https://doi.org/10.1016/s0092-8674(00)80700-x

Schürholz, A.-K., López-Salmerón, V., Li, Z., Forner, J., Wenzl, C., Gaillochet, C., Augustin, S., Barro, A.V., Fuchs, M., Gebert, M., Lohmann, J.U., Greb, T., Wolf, S., 2018. A Comprehensive Toolkit for Inducible, Cell Type-Specific Gene Expression in Arabidopsis. Plant Physiol. 178, 40–53. https://doi.org/10.1104/pp.18.00463

Serrano-Mislata, A., Schiessl, K., Sablowski, R., 2015. Active Control of Cell Size Generates Spatial Detail during Plant Organogenesis. Curr. Biol. CB 25, 2991–2996. https://doi.org/10.1016/j.cub.2015.10.008

StarDist - Object Detection with Star-convex Shapes, 2023.

Tran, A.P., Yan, S., Fang, Q., 2020. Improving model-based functional near-infrared spectroscopy analysis using mesh-based anatomical and light-transport models. Neurophotonics 7, 015008. https://doi.org/10.1117/1.NPh.7.1.015008

Wang, Y., Shirakawa, M., Ito, T., 2023. Arrest, Senescence and Death of Shoot Apical Stem Cells in Arabidopsis thaliana. Plant Cell Physiol. 64, 284–290. https://doi.org/10.1093/pcp/pcac155

Werner, T., Holst, K., Pörs, Y., Guivarc’h, A., Mustroph, A., Chriqui, D., Grimm, B., Schmülling, T., 2008. Cytokinin deficiency causes distinct changes of sink and source parameters in tobacco shoots and roots. J. Exp. Bot. 59, 2659–2672. https://doi.org/10.1093/jxb/ern134

Wolny, A., Cerrone, L., Vijayan, A., Tofanelli, R., Barro, A.V., Louveaux, M., Wenzl, C., Strauss, S., Wilson-Sánchez, D., Lymbouridou, R., Steigleder, S.S., Pape, C., Bailoni, A., Duran-Nebreda, S., Bassel, G.W., Lohmann, J.U., Tsiantis, M., Hamprecht, F.A., Schneitz, K., Maizel, A., Kreshuk, A., 2020. Accurate and versatile 3D segmentation of plant tissues at cellular resolution. eLife 9, e57613. https://doi.org/10.7554/eLife.57613

Yadav, R.K., Perales, M., Gruel, J., Girke, T., Jönsson, H., Reddy, G.V., 2011. WUSCHEL protein movement mediates stem cell homeostasis in the Arabidopsis shoot apex. Genes Dev. 25, 2025–2030. https://doi.org/10.1101/gad.17258511

Zhou, Y., Liu, X., Engstrom, E.M., Nimchuk, Z.L., Pruneda-Paz, J.L., Tarr, P.T., Yan, A., Kay, S.A., Meyerowitz, E.M., 2015. Control of plant stem cell function by conserved interacting transcriptional regulators. Nature 517, 377–380. https://doi.org/10.1038/nature13853

Zhou, Y., Yan, A., Han, H., Li, T., Geng, Y., Liu, X., Meyerowitz, E.M., 2018. HAIRY MERISTEM with WUSCHEL confines CLAVATA3 expression to the outer apical meristem layers. Science 361, 502–506. https://doi.org/10.1126/science.aar8638

